# Forest-weighted non-linear causal inference for high-dimensional and multimodal time series

**DOI:** 10.64898/2026.09.24.754251

**Authors:** Shigeki Kishi

## Abstract

Forecasting and causal inference in complex, non-linear dynamical systems are fundamentally challenging due to noisy and high-dimensional time-series data. While Empirical Dynamic Modeling addresses this via state-space reconstruction in Euclidean space, it suffers from the “curse of dimensionality,” distorting distance measurements as dimensions increase. This study proposes Forest-Weighted S-map (FORWS) and Forest-Weighted Causal Inference (FORWC), utilizing adaptive “forest weights” derived from random forest ensembles instead of Euclidean metrics. Using simulated and empirical ecological datasets, I demonstrate that these methods exhibit comparable or improved forecasting skill relative to conventional tools, while offering substantial resilience against dynamic process noise and mitigating the curse of dimensionality. Notably, FORWC enables multimodal causal inference, successfully extracting directed interactions between high-dimensional acoustic vectors and scalar temperature data monitored in a honeybee hive. By bridging chaotic physics with machine learning, this framework accurately extracts intrinsic manifold topology, providing a robust paradigm for uncovering causal networks in complex real-world environments.

## Introduction

From natural environments to biological signals, real-world systems are increasingly recognized as deterministic chaotic systems characterized by state-dependent interactions. In such nonlinear dynamics, traditional parametric statistical methods often struggle to capture true causal relationships, as they typically assume that the governing parameters remain constant over time. To overcome this limitation, Empirical Dynamic Modeling (EDM) has been developed to reconstruct the system’s attractor manifold directly from observed time-series data(Munch et al., 2023). Grounded in Takens’ delay embedding theorem(Takens, 1981), EDM has achieved significant success in forecasting complex dynamics and detecting causality through methods such as S-map(Sugihara, 1994) and Convergent Cross Mapping (CCM)(Sugihara et al., 2012), which operate without requiring prior knowledge of the underlying governing equations.

However, recent advances in observation technologies, such as the Internet of Things (IoT), have generated an unprecedented volume of ultra-high-dimensional datasets, confronting EDM with a significant mathematical barrier. Conventional S-map and CCM frameworks rely entirely on isotropic Euclidean distances when evaluating the proximity between a target state and its neighbors on the reconstructed manifold. Because Euclidean distance is calculated uniformly across all dimensions, the accumulation of small fluctuations in irrelevant variables, or observational noise, obscures true proximity in high-dimensional embedded spaces(Ye & Sugihara, 2016). Due to this “curse of dimensionality,” dynamically similar states are often pushed far apart, rendering appropriate local weighting impossible. Multiview Embedding approach (MVE)(Ye & Sugihara, 2016) and Multiview Distance Regularized S-map (MDR S-map)(Chang et al., 2021) attempted to overcome this by ensembling distance information across low-dimensional subspaces, but proximity measurements based on Euclidean metrics remain problematic because local manifolds in the state space are frequently distorted in a non-Euclidean manner(Kawatsu, 2024).

Adaptive sample weighting derived from machine learning algorithms offers a potential breakthrough for this constraint(Lin & Jeon, 2006). Algorithms such as k-nearest neighbors, support vector classifiers, random forests, and neural networks all process information by evaluating the proximity between samples in a feature space, though their internal definitions of similarity differ(Hastie et al., 2009). For instance, in random forests, sample similarity is measured by the co-occurrence frequency of samples within the same terminal nodes (leaves) across an ensemble of decision trees, rather than by geometric Euclidean distance(Breiman, 2001; Wager & Athey, 2018). Consequently, random forests inherently function as an adaptively weighted nearest-neighbor method that partitions the feature space according to variable importance(Lin & Jeon, 2006). Building on this, the Generalized Random Forest (GRF) framework(Athey et al., 2019) introduced ‘forest weights’ derived from an adaptive weighting function. This function determines the weights based on the frequency of samples falling into the same terminal nodes, effectively accounting for the local density and structure of the data. These weights can serve as a metric that reflects the intrinsic structure of the data manifold more accurately than conventional Euclidean distance measures.

This study proposes Forest-Weighted S-map, FORWS, and Forest-Weighted Convergent Cross Mapping, FORWC. These methods integrate deterministic chaotic physics with the optimization theory of machine learning. Their most defining feature is the use of an adaptive forest weight matrix rather than an isotropic Euclidean distance matrix (Fig. 1).

**Fig. 1:**
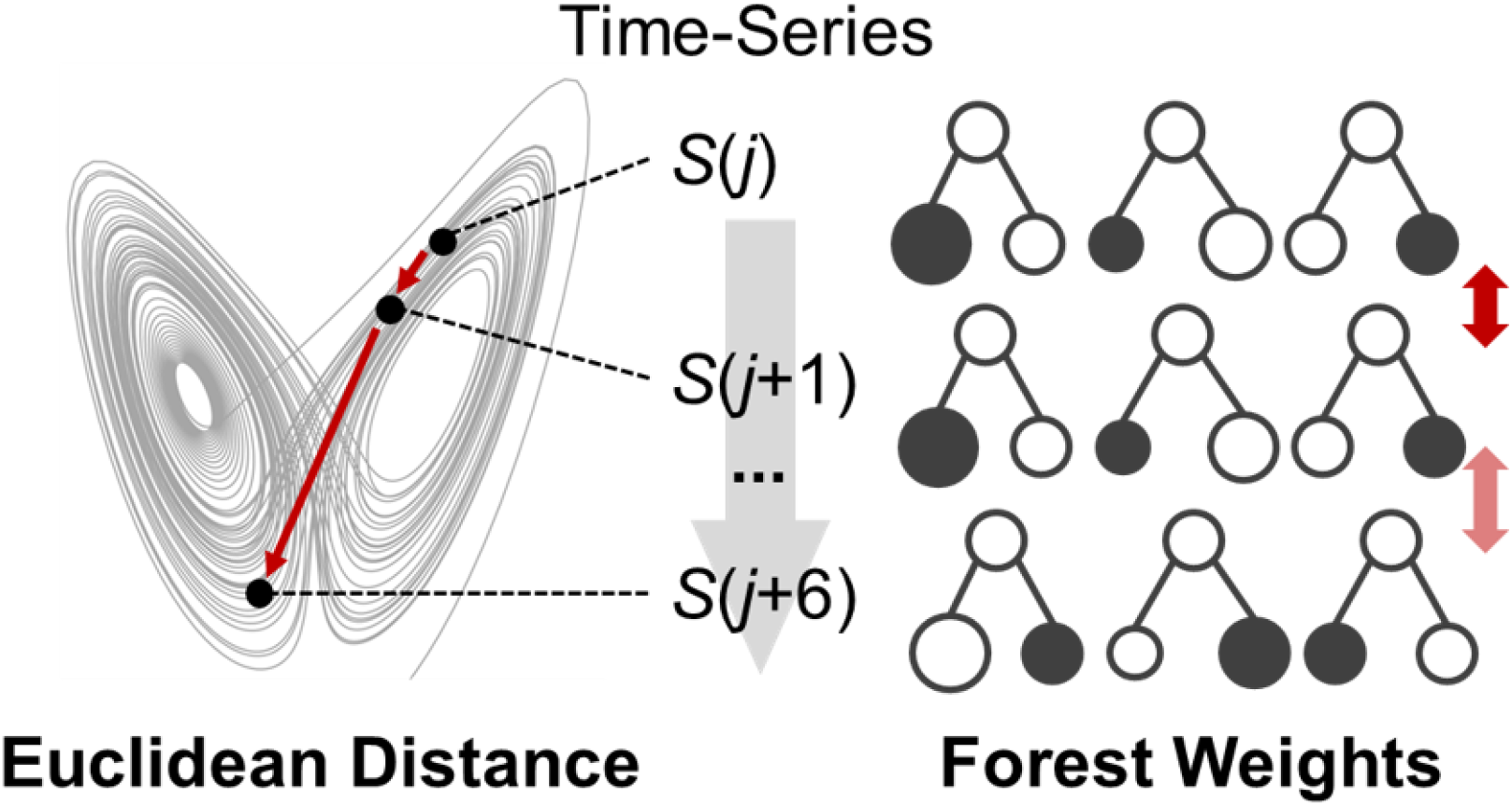
Conceptual comparison of state proximity metrics. Conventionally, Euclidean distance (red arrows) has been used to measure the proximity between historical states embedded in a multidimensional state space (left). In contrast, this study introduces forest weights, quantifying state similarity based on the frequency of samples sharing the same terminal leaves across member decision trees (right).

I developed analytical functions for FORWS, FORWC, and associated preprocessing tools to determine the optimal embedding dimensions and effective time lags in R. First, I evaluated the methods using simulated three-species interaction dynamics. Second, I applied the framework to empirical population dynamics of *Paramecium* and *Didinium* to compare the predictive accuracy of FORWS against MDR S-map. Finally, I demonstrated multimodal and high-dimensional causal inference using acoustic and temperature time-series recorded in a honeybee hive.

## Results

### Causal inference in chaotic dynamics and noisy environments

I simulated the population dynamics of three interacting species (*X, Y*, and *Z*) using discrete-time difference equations for 300 steps (see Methods). In the formula, *X* and *Y* exert negative effects on *Y* and *Z*, respectively (Fig. 2a).

**Fig. 2:**
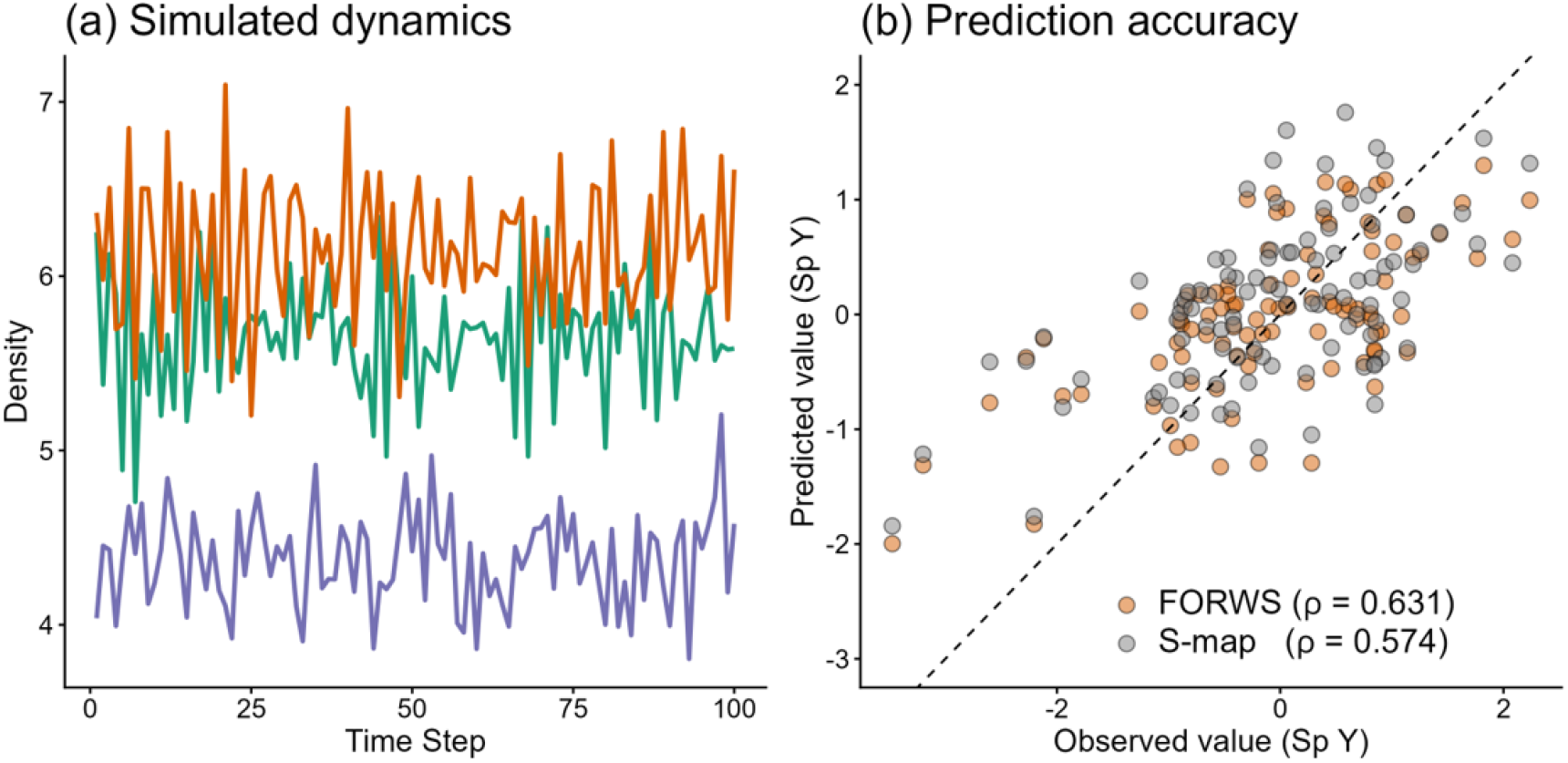
Performance comparison of FORWS and conventional S-map using simulated multispecies dynamics. (a) Time series of the simulated densities for three interacting species (*X*_*t*_, *Y*_*t*_, and *Z*_*t*_). The plot displays the initial 100 time steps out of a total of 300 steps. (b) Comparison of observed versus predicted values for species Y. Orange and grey circles represent predictions generated by Forest-Weighted S-map (FORWS) and conventional S-map, respectively. The dashed line indicates a perfect 1:1 prediction.

I used the latter 200 steps for analyses. The optimal embedding dimension was 1 for all species (*E\**_*X*_ = *E\**_*Y*_ = *E\**_*Z*_ = 1) and the effective time-delay was 1. To compare prediction accuracy between conventional S-map and FORWS, Pearson correlation coefficient, *ρ*, between simulated and predicted values of *Y*_*t+1*_ was calculated when *Y*_*t+1*_ was predicted by *X*_*t*_ and *Y*_*t*_, *Y*_*t-1*_, *Y*_*t-2*_ in test data (the latter half of the data) on the manifold built by train data (the former half). The *ρ* value was 0.631 for FORWS, where minimal node size was 3 and the ridge regression parameter was *λ* = 1e-6, and 0.574 for S-map, where the transforming parameter was *θ* = 0.75.

FORWC only detected two true causalities (*X* to *Y, Y* to *Z*, where “*X* to *Y*” is supported by the successful cross-mapping (reconstruction) from *Y* to *X*) among six directions with no misdetections (Fig. 3a), while CCM detected a false-positive causality (*X* to *Z*) and a marginal (*Y* to *X*) other than true causalities (Fig. 3b).

**Fig. 3:**
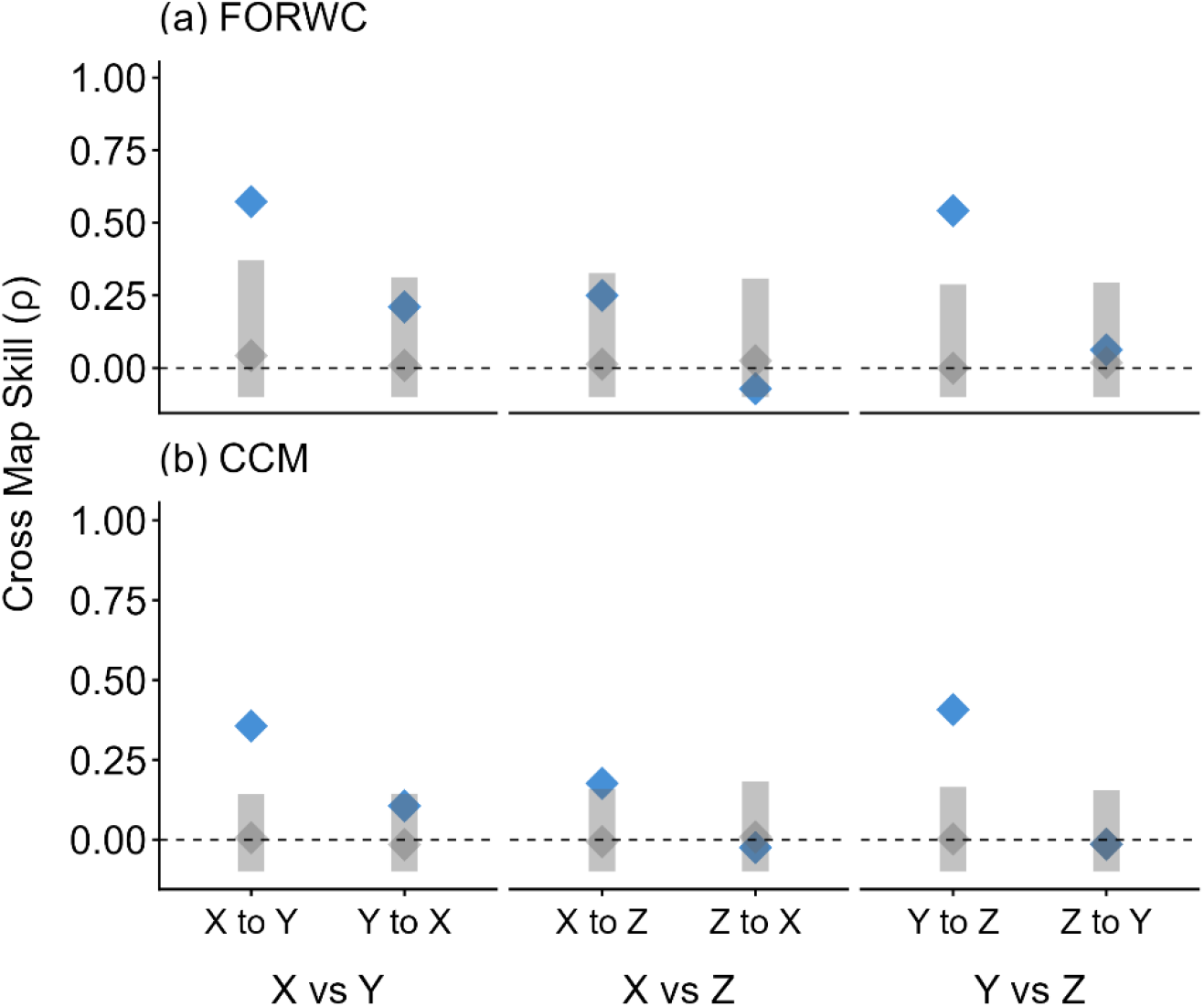
Comparison of causality inference between FORWC and conventional CCM. Cross-map skill (*ρ*) obtained using (a) Forest-Weighted Convergent Cross Mapping (FORWC) and (b) conventional CCM for simulated multispecies dynamics. Blue diamonds represent the mean cross-map skill of the observed data. Grey diamonds and shaded bars indicate the mean and 95% confidence intervals, respectively, derived from surrogate target data. Time-series causality is considered significant when the observed skill (blue diamond) exceeds the upper bound of the surrogate distribution (grey bar).

To rigorously evaluate the robustness of these methods against dynamic process noise, I systematically varied the noise strength in the three-species system from 0.02 to 0.16 and assessed the causality from *X* to *Y* (Fig. 4).

**Fig. 4:**
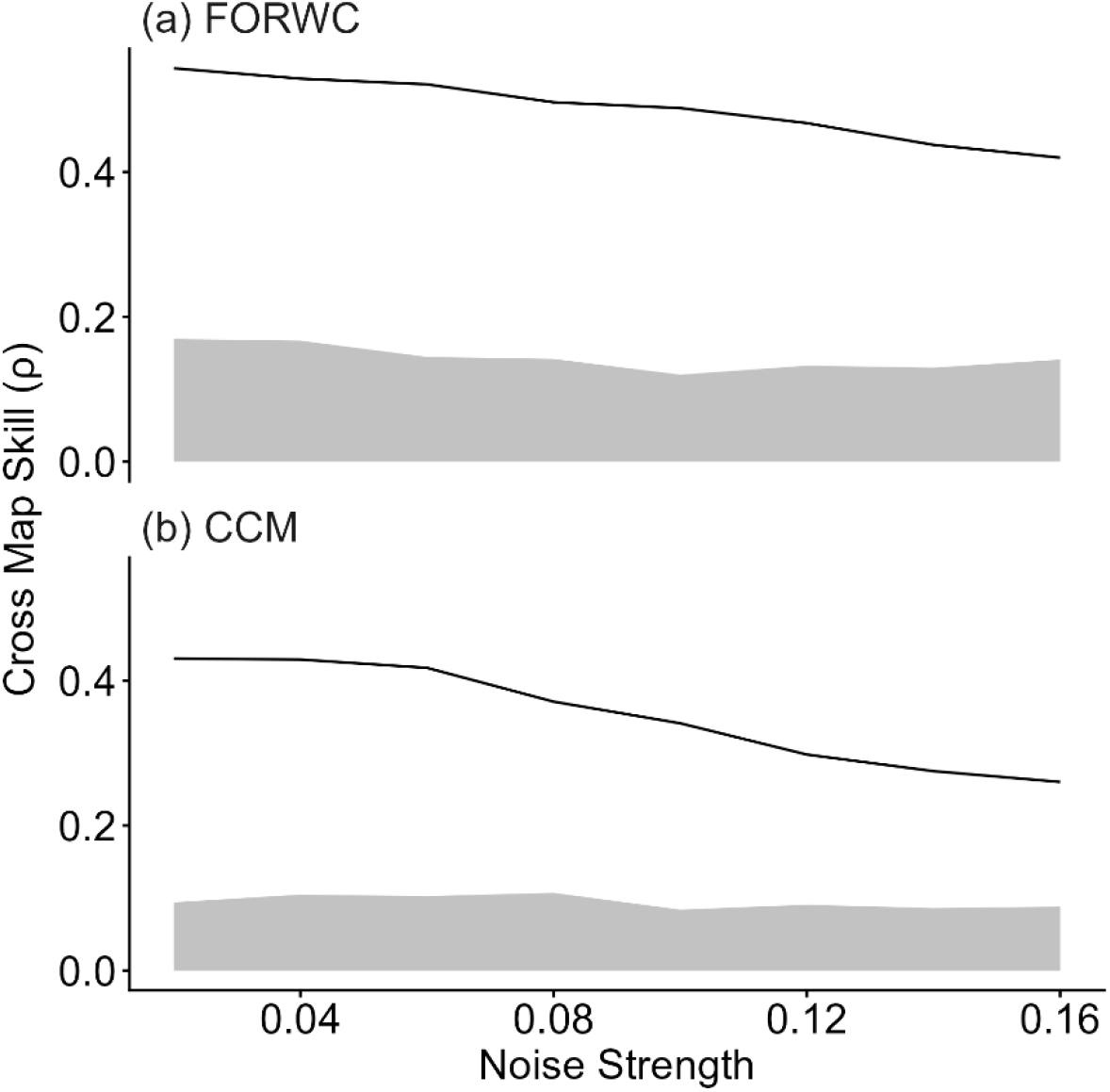
Cross-map skill (ρ) from Y to X (causal direction X to Y) at the library size 100 and test size 100 was shown when noise strength was changed from 0.02 to 0.16, for (a) Forest-Weighted Convergent Cross Mapping (FORWC) and for (b) conventional CCM. Solid lines represent the mean cross-map skill of the observed data. Grey zones indicate 95% confidence intervals, derived from surrogated target data. Time-series causality is considered significant when the observed skill (solid line) exceeds the upper bound of the surrogate distribution (grey zone).

While both FORWC and conventional CCM successfully detected significant causality (observed skill exceeding the 95% surrogate confidence intervals) across all noise levels, their predictive performance trajectories diverged markedly. As noise increased, the cross-map skill of conventional CCM degraded rapidly, dropping from approximately 0.45 to 0.25 (Fig. 4b). This vulnerability arises because the isotropic Euclidean distance struggles to distinguish true manifold geometry from state-space distortions caused by process noise. In contrast, FORWC maintained a remarkably high and stable cross-map skill, retaining more than 0.4 even at the maximum noise level (Fig. 4a). By optimizing the minimum terminal node size to prevent overfitting (10 in this case), the random forest ensemble effectively acts as an adaptive structural regularizer. This allows FORWC to smooth out stochastic fluctuations while accurately capturing the intrinsic causal signal, proving its resilience in highly noisy environments.

### Nonlinear phase shifts in empirical predator-prey dynamics

Empirical population dynamics of two plankton species, the predator *Didinium nasutum* (Dn) and its prey *Paramecium aurelia* (Pa), were analyzed using FORWS and MDR S-map (Kawatsu, 2024; Sugihara et al., 2012). The bacterium *Aerobacter aerogenes* was provided as food resource for Pa. The dataset consists of three experiments in which Cerophyl, powder of dried young wheat leaves used as a bacterial growth medium, concentration was intermediate (0.500 g/L) in experiment 1 and 3, and low (0.375 g/L) in experiment 2. In experiment 3, a researcher removed a constant ratio of the predator from an arena at a constant interval(Jost & Ellner, 2000; Veilleux, 1979).

Estimated effects of Pa on Dn, and of Dn on Pa were compared between FORWS and MDR S-map, a state-of-the-art Euclidean method designed to mitigate the curse of dimensionality (Fig. 5ab).

**Fig. 5.**
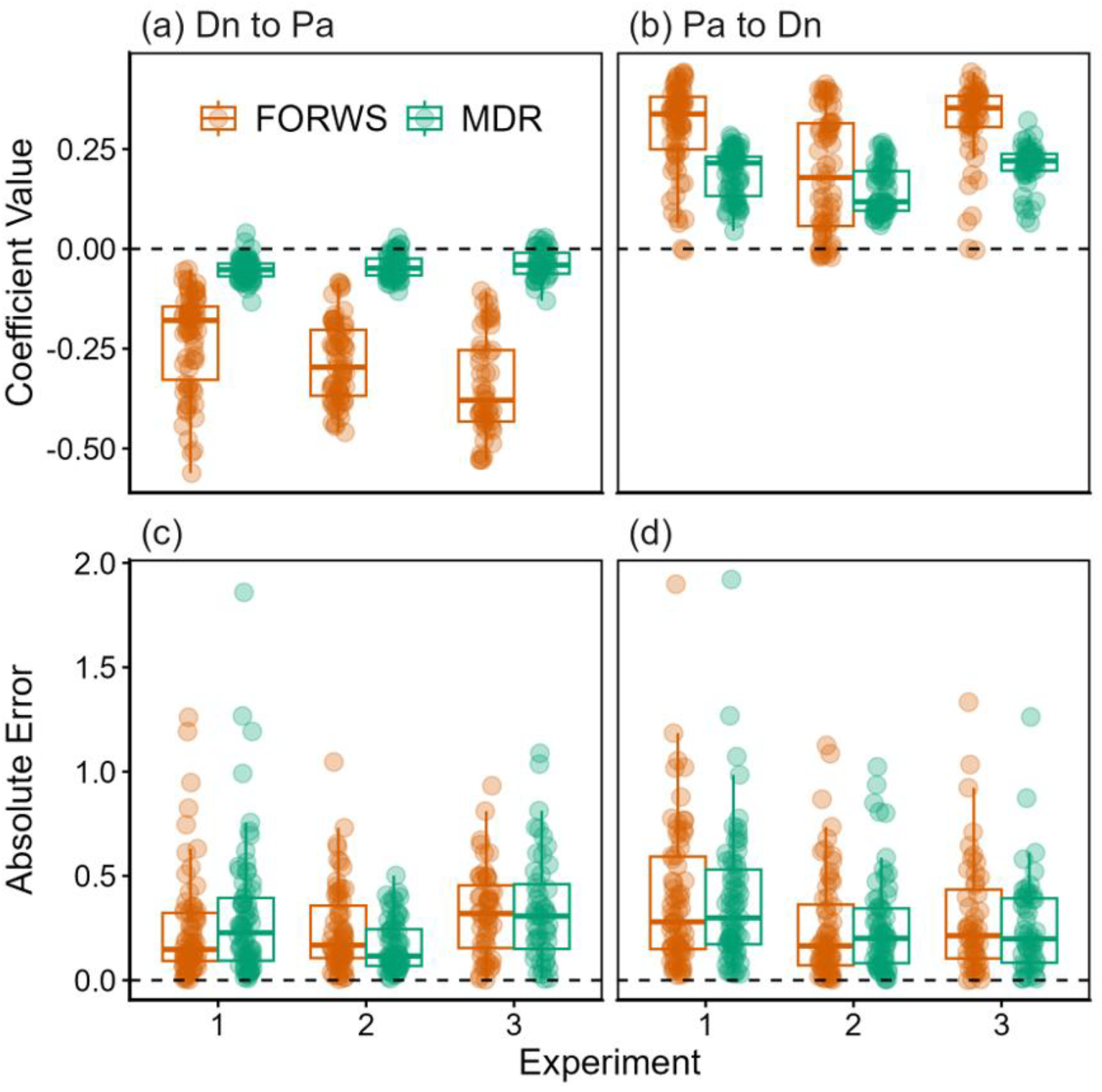
Estimated interaction strengths and prediction accuracies for the *Didinium* (predator, Dn)–*Paramecium* (prey, Pa) dynamics. Local linear coefficients quantifying the effect of (a) *Didinium* (*Dn*_*t*_) on Paramecium (*Pa*_*t*_) and (b) Paramecium (*Pa*_*t-2*_) on *Didinium* (*Dn*_*t*_) across three experimental conditions (Exp. 1: medium nutrition; Exp. 2: low nutrition; Exp. 3: medium nutrition with proportional predator removal). Absolute prediction errors (differences between observed and predicted values) corresponding to the models for (c) Dn to Pa and (d) Pa to Dn. Estimations derived from FORWS and MDR S-map are indicated by orange and green boxplots with jittered points, respectively. In the boxplots, the upper, center, and bottom lines indicate the first, second (median), and third quartiles respectively, and the whiskers extend to 1.5 interquartile range. The dashed horizontal line represents zero in all panels (a–d).

The optimal number of embedding dimensions was *E*\* = 2 for both species, and the time-delay was 1 for both directions. Then *Pa*_*t*_, *Pa*_*t-1*_ and *Dn*_*t*_ were used to predict *Pa*_*t+1*_, while *Dn*_*t*_, *Dn*_*t-1*_ and *Pa*_*t*_ were used to predict *Dn*_*t+1*_. To evaluate this statistically, I employed a linear mixed model. The coefficient value of the interacting species was set as the response variable, while the method, causal direction, and their interaction were included as explanatory variables, with the experiment identity as a random effect. This model detected significant difference between MDR S-map and FORWS (*t* = 12.7, *p* < 0.0001) and between directions (*t* = 35.41, *p* < 0.0001) and the interaction (*t* = –6.61, *p* < 0.0001). The local linear coefficient values, representing effecf of *Dn*_*t*_ on *Pa*_*t+1*_, were predominantly negative while those for the opposite direction were positive (Fig. 5ab). Coefficient values of *Dn*_*t*_ on *Pa*_*t+1*_ estimated by FORWS were more negative than MDR S-map (*p* < 0.0001, Fig. 5a). Coefficient values of *Pa*_*t*_ on *Dn*_*t+1*_ estimated by MDR S-map were larger than FORWS (*p* = 0.0013, Fig. 5b). Meanwhile a generalized linear mixed model, in which response variable was absolute residual value with gamma distribution, explanatory variables were method and causal direction and the interaction, and the random effect was experiments, detected no significant differences of absolute residual values between FORWS and MDR S-map (*t* = –0.62, *p* = 0.54) and between causal directions (*t* = –1.78, *p* = 0.075) and the interaction effect (*t* = 0.35, *p* = 0.73) (Fig. 5cd).

### Multimodal causal inference uncovers asymmetric thermoregulatory dynamics

Using a multimodal time-series dataset comprising internal hive temperature and simultaneous 12-second acoustic recordings sampled at 15-minute intervals, I examined the causality between these two modalities. Ecologically, bidirectional causality is highly plausible: ambient temperature influences the bees’ acoustic behavior (e.g., buzzing), and the mechanical energy from this behavior, in turn, affects the hive’s temperature. Each raw audio recording (an example shown in Fig. 6a), represented by buzzing sound in a beehive, was processed into non-overlapping 2-second segments.

**Fig. 6:**
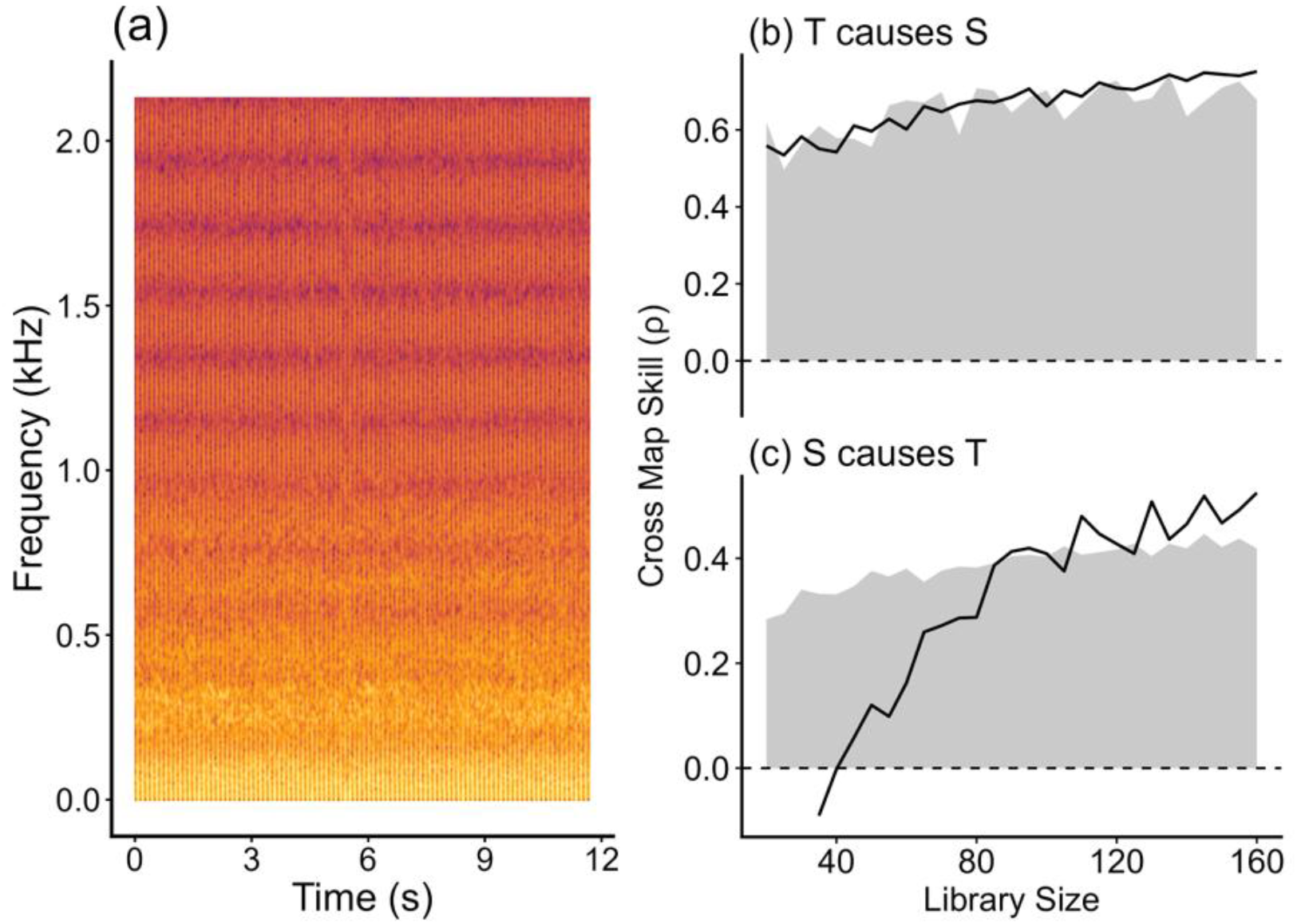
Bidirectional causal inference between internal hive temperature and the acoustic environment. (a) A representative spectrogram of a 12-second acoustic recording sampled at 15-minute intervals. Color intensity, ranging from dark to yellow, represents acoustic amplitude. (b, c) Convergent cross mapping (CCM) using FORWC to test the causality (b) from temperature (T) to sound (S) (cross-mapping from sound to temperature) (c) from sound (S) to temperature (T) (cross-mapping from temperature to sound). Solid black lines indicate the cross-map skill (Pearson correlation coefficient, *ρ*) derived from the observed data. The grey shaded regions represent the 95% prediction intervals of the null distribution derived from surrogate data.

All segments from a single record were averaged to a representative segment. The segment matrix was then partitioned into a 10 × 10 grid (frequency by time), and the median values within each bin were vectorized into a 100-dimensional feature vector.

First, causalities between each of the 100 time series of the acoustic time-series and temperature were examined using FORWC. Forty-nine acoustic time-series were estimated as causal for the directions from sound to temperature, while temperature was causal for 61 acoustic time-series. The optimal time lags were 0 and −3 for the direction from temperature to sound and from sound to temperature, respectively. Using these conditions, FORWC was carried out for both directions. In both directions, cross map skill between observed data went up beyond the 95% prediction interval of that between observed and surrogate data when the library size increased (Fig. 6bc). For the causal direction from temperature to sound, the cross-map skill consistently exceeded the upper bound of the 95% surrogate confidence interval (Fig. 6b).

Meanwhile, the cross map skill from temperature to sound slowly increased with library size (Fig. 6c).

## Discussion

This study demonstrates that S-map and convergent cross mapping (CCM) based on forest weights derived from random forests, rather than Euclidean distances, function effectively and perform comparably to, or slightly outperform conventional measures across both simulated and real-world datasets. By replacing the isotropic Euclidean kernel with the adaptive weighting function of Generalized Random Forests (GRF), I established FORWS and FORWC—a robust, data-driven framework that bridges deterministic chaotic physics with machine learning optimization. Remarkably, this framework enables multimodal causal inference directly between high-dimensional vectors and scalar time series, as demonstrated by the interactions between honeybee acoustic features and internal hive temperature. By effectively overcoming the “curse of dimensionality”, this approach provides a scalable and versatile solution for causal discovery in complex, real-world ecosystems.

The comparative performance between simulated and experimental datasets provides critical insights into the mechanism of FORWS. In the low-dimensional simulated dataset (Fig. 2), FORWS showed comparable or slightly superior performance over S-map. This superiority stems from the adaptive nature of forest weights; while the standard S-map relies on a fixed exponential decay kernel 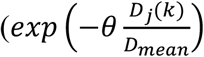, where a coefficient value is *θ*, the distance between sample *j* and *k* is *D*_*j*_(*k*), and the mean distance is *D*_*mean*_)(Sugihara, 1994), FORWS directly calculates the sample weights, optimizing the local neighborhood through the tuning of regularization parameter (*λ*) in weighted multiple regression and the minimum terminal node size. Once tuned with these hyper parameters, the model captured the local curvature of the manifold, as accurately as, or better than, an isotropic Euclidean distance, even in artificial deterministic systems, effectively avoiding the overestimation of noise.

Furthermore, the comparative analysis of the simulated three-species dynamics (Fig. 3) highlights a critical advantage of FORWC in mitigating false-positive causal detections. In deterministic dynamical systems, conventional CCM often struggles with generalized synchronization or transitive indirect causal chains (e.g., incorrectly identifying causality from *X* to *Z*). This vulnerability arises because isotropic Euclidean distances can be easily deceived by globally correlated trajectories. In contrast, FORWC successfully isolated only the true, direct causal interactions (*X* to *Y* and *Y* to *Z*). By constructing neighborhoods based on the frequency of terminal node co-occurrence, the random forest ensemble acts as an adaptive feature-space partitioner that implicitly weights the most informative dimensions. This allows FORWC to filter out spurious correlations and distinguish the true causal manifold topology from generalized synchronization, thereby preventing the overestimation of causality that plagues traditional distance-based methods.

Comparable prediction accuracy was also observed in analyzing the experimental population dynamics of *Didinium* and *Paramecium* (Fig. 5). The fact that FORWS achieved absolute prediction errors equivalent to MDR S-map—which relies on complex low-dimensional ensembling to measure distance—demonstrates that the forest weight approach is inherently robust against noise and the curse of dimensionality. These results suggest that forest weights can serve as a highly precise estimator of state-space transitions, giving high prediction accuracy to FORWS.

Although the overall prediction accuracy did not differ between the two methods for the experimental datasets, the estimated local linear coefficient values exhibited distinct distributions. A larger variance of the coefficient values estimated by FORWS may suggest a more sensitive localization in the state-space. Particularly, the negative coefficient values of Pa on Dn in Experiment 2 reflect the delayed crash of the predator (Dn) following the overexploitation of the prey (Pa). In other words, FORWS successfully predicted the critical phase shift where an increase in prey subsequently leads to a sharp decline in predators due to resource exhaustion. This suggests that FORWS can be a powerful next-generation tool to deeply understand complex ecological systems with highly non-linear phase transitions.

The most characteristic feature of FORWC is its capability to perform multimodal causal inference directly on multidimensional data. In the honeybee hive analysis, FORWC detected bidirectional causalities between a 100-dimensional acoustic vector and a one-dimensional scalar temperature (Fig. 6ab). Notably, it uncovered a clear asymmetry in the time lags: while the honeybees’ acoustic behavior responded immediately to temperature changes (*tp* = 0), the collective acoustic energy (e.g., shivering) required a physical latency of approximately 45 minutes (*tp* = –3) for its generated mechanical energy to meaningfully elevate the internal hive temperature.

As observational technologies such as IoT sensors, camera traps, and environmental DNA (eDNA) become ubiquitous, researchers are increasingly overwhelmed by high-dimensional, multimodal datasets. Because FORWC processes similarity through terminal node co-occurrence rather than geometric distance, this framework is theoretically expandable to track causalities between continuous image streams or hierarchical ecological networks. Furthermore, the core concept of adaptive weighting can be extended beyond random forests to other advanced machine learning architectures, such as gradient boosting algorithms, to further optimize state-space reconstruction. FORWS and FORWC provide a completely model-free, biologically interpretable method to uncover the hidden causal networks driving these complex systems, opening a new frontier for nonlinear time-series analysis in ecology and beyond.

## Materials and Methods

### Formulation of Forest Weights and Local Linear Regression

Forest weights are derived from the Generalized Random Forests (GRF) framework (Athey et al., 2019). Let *X*_*i*_ be the reconstructed state-space vector (or high-dimensional feature vector) at time *i*, and *Y*_*i*_ be the corresponding target value. A regression forest consisting of *B* decision trees (e.g., *B* = 2000) is trained on the state-space. For a given query state *x*, the forest weight *α*_*i*_(*x*) assigned to the *i*-th training observation is defined by the frequency with which *X*_*i*_ falls into the same terminal node (leaf) as the query *x* across all *B* trees:

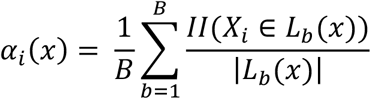

where *L*_*b*_(*x*) denotes the set of training samples fallen in the same leaf with *x* in the *b*-th tree, |*L*_*b*_(*x*)| is the number of samples in that leaf, and II(·) is the indicator function, which returns 1 if *X*_*i*_ is found with *x* in a node of the *b*-th tree. Then forest weight not only evaluates whether a training sample *X*_*i*_ and the query *x* fall into the same terminal node, but also scales the similarity by the density (the number of samples) of that node. A smaller number of samples in the shared leaf (|*L*_*b*_(*x*)|) indicates that the two states occupy a highly specific, localized region of the state space, thereby assigning them a heavier weight. The final forest weight, *α*_*i*_(*x*), is then computed as the ensemble mean of these similarity values across all *B* decision trees. This property allows the algorithm to dynamically adjust the resolution of the local neighborhood based on the underlying complexity of the manifold.

In generating these weights, I optimized the minimum node size of the trees rather than the maximum tree depth. This hyperparameter acts as a structural regularizing factor by enforcing a minimum number of samples in each leaf, guaranteeing a stable minimum neighborhood size. This explicit restriction prevents the isolation of samples on the edges of the manifold and ensures to estimate the proper forest weight matrix including those outlying samples. Subsequently, the prediction for the target variable *Ŷ*(*x*) is estimated through a locally weighted linear regression using the derived forest weights. To address potential multicollinearity among high-dimensional features and to stabilize the inverse matrix computation, I incorporated a ridge (L2) regularization penalty, *λ*. The local linear coefficients 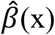 are computed by minimizing the penalized weighted least squares:

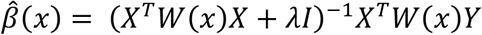

where *W*(*x*) is a diagonal matrix containing the forest weights *α*_*i*_(*x*), and *I* is the identity matrix. The final prediction is given by

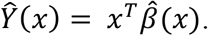

To identify the optimal model configuration, I conducted a grid search across a range of values for the minimum node size and the Ridge penalty, *λ*. The final parameter set was selected based on the minimization of the root mean square error (RMSE) through cross-validation, ensuring that the model achieved the highest predictive stability without over-fitting to observational noise.

### Analysis Procedure of FORWC and FORWS

#### Step 1. Estimation of Optimal Embedding Dimension

The optimal embedding dimension *E\** is fundamental for reconstructing the system’s manifold. For univariate scalar data such as temperature, *E\** was identified by evaluating the prediction skill (Pearson correlation coefficient: *ρ*) and RMSE based on regression forest across various dimensions. For high-dimensional acoustic data, I utilized a multi-regression forest approach to determine the *E\** that best captures the multidimensional state-space structure, ensuring that the complex dynamics are adequately captured. In the default settings of the function, the values of *ρ* and RMSE were calculated from *E* = 0 to *E* = 10, and the ratios of train and test data were 0.8 and 0.2, respectively, randomly sampled from original data.

#### Step 2: Determination of Optimal Time Lag

To account for the inherent time-lagged effect of causal time-series on recipient time-series, I explored the optimal time lag, *tp*. By evaluating the cross-map skill over a defined range of *tp*, I identified the lag at which the predictive information transfer peaked. This step is critical for prediction and causal inference; a significant peak at *tp* < 0 (negative lag) provides evidence that the recipient time-series contains past information about the causal time-series, consistent with physical precedence. In the default settings of the function, the values of *ρ* and RMSE were calculated from *tp* = –5 to *tp* = 5, using the fixed *E\**.

#### Step 3: FORWS for Prediction and FORWC for Causal Inference

Using the optimized *E, tp*, and hyperparameters (*λ*, minimal node size), I executed the final models: Forest-Weighted S-map (FORWS) for time-series forecasting and Forest-Weighted Causal Inference (FORWC) for causality detection.

FORWS forecasts the future states of a target time series by leveraging both its own historical and other relevant variables. These variables are embedded into a multidimensional state space where sample similarities are quantified via the forest weight matrix. Unlike the conventional S-map, which relies on a fixed Euclidean distance kernel to assign weights, FORWS adaptively evaluates the sample distribution within the state space based on the recursive multidimensional partitioning of the random forest. These forest weights are subsequently used in a regularized locally weighted linear regression to estimate the response variable, facilitating highly accurate and noise-robust forecasting of complex nonlinear dynamics. In the default settings of the function, the number of decision trees was 2000, the minimal sample size in a node was 5, and the regulator factor, *λ*, was 0.0001.

FORWC also follows the logic of the conventional CCM. If the recipient time series can accurately reconstruct—or “predict”—the causal time series, often in the past, better than null models, it means that information about the causal time series is embedded in the recipient time series. For causality test, it is examined whether this “cross-map skill” increases with the library size of the recipient data and eventually converges beyond 95% confidence interval of null models. FORWC uses forest weights, instead of Euclidean distances in CCM, to evaluate the sample distribution in the recipient manifold. In the default settings of the function, the number of decision trees was 2000, the minimal sample size in a node was 5, and the regulator factor, λ, was 0.0001.

Null data of the causal time-series were made by the Ebisuzaki surrogate method(Ebisuzaki, 1997). This approach generates surrogate time series that preserve the power spectrum of the original data while randomizing the phases. I constructed a null distribution of cross-map skills from 50 surrogates to calculate 95% confidence intervals. Causality was deemed statistically significant only when the observed skill consistently exceeded the upper value of 95% confidence intervals. To prevent any information leakage, “cheating”, from the true causal structure into the null model, the random forest and its corresponding forest weights were recalculated from scratch for each surrogate time series. This ensures a strictly independent null distribution for statistical testing.

### Simulated 3 species competition dynamics

To evaluate the basic performance of FORWS and FORWC, I utilized a artificial dataset generated from a chaotic three-species discrete-time competition model.

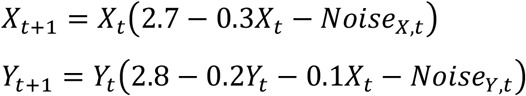

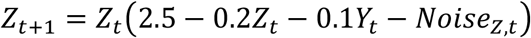

The densities of species *X, Y* and *Z* are *X*_*t*_, *Y*_*t*_, and *Z*_*t*_, respectively. Two causal interactions were set from *X*_*t*_ to *Y*_*t*_, and from *Y*_*t*_ to *Z*_*t*_. Initial densities (*X*_*1*_, *Y*_*1*_, *Z*_*1*_) are 1.0. The system was simulated for 300 time steps, and the first 100 steps were discarded (Fig. 1a). The value of *Noise*_*sp,t*_ was randomly drawn from a Gaussian distribution with the mean value, 0, and standard deviation, 0.1.

When I explored the optimal number of embedding dimensions, *E\**_*sp*_, for each species by the function written above, that was 1 (*E\**_*x*_ = *E\**_*y*_ = *E\**_*z*_ = 1). The optimal time lag was 1 for all directions following the formula. Then *Y*_*t+1*_ was predicted by *X*_*t*_ and *Y*_*t*_ by using FORWS and S-map. In FORWS, the best values of *λ* and the minimal sample size were 1e–6 and 3, respectively. In S-map, the best value of *θ* was 0.75. Causality was tested for all 6 directions by using FORWC and CCM.

To evaluate the robustness against dynamic process noise, the standard deviation of the noise, *Noise*_*sp,t*_ was incrementally increased from 0.02 to 0.16. To ensure a strictly fair comparison between the two methods, both FORWC and CCM were evaluated using a library size and test size of 150 and 50, respectively, with cross-map skills averaged over 20 random sampling iterations. For FORWC, the minimum terminal node size was set to 10. This explicit restriction acts as a structural regularizer, preventing the decision trees from overfitting to the dynamic process noise while extracting the true causal neighborhood.

### Experimental dynamics of predator and prey

To compare prediction performance between FORWS and MDR S-map on empirical data, I utilized the classic predator-prey time series from the microcosm experiments conducted by Veilleux (Veilleux, 1979) and subsequently digitized and analyzed by following studies (Jost & Ellner, 2000; Kawatsu, 2024; Sugihara et al., 2012). This dataset provides a high-resolution record of the population dynamics between the predatory ciliate *Didinium nasutum* and its prey *Paramecium aurelia*. I specifically focused on three datasets that differ in experimental settings: (1) a medium nutrient condition with Cerophyl (powder of dried young wheat leaves given for the bacterium) concentration (0.500 g/L), (2) a poor nutrition condition with low Cerophyl concentration (0.375 g/L), and (3) a medium nutrient condition, identical with (1), but subjected to regular, artificial removal of the predator. In this third condition, a specific proportion of the *Didinium* population was manually removed during the regular transfer of cultures to fresh medium, simulating a density-independent mortality event on the higher trophic level.

For the *Didinium*–*Paramecium* time series, I first estimated the optimal embedding dimension (*E\**) and time lag (*tp*) using the grid search procedure as described above. Once the optimal manifold was reconstructed, I applied both FORWS and the MDR S-map to predict the population dynamics and extract the local linear coefficients at each time step. To evaluate prediction accuracy, I calculated absolute differences between observed and the predicted values. In predator-prey systems, a globally positive coefficient typically indicates a synchronized trend (i.e., higher prey abundance correlates with higher predator abundance).

### Multimodal causal inference

As a representative example of complex, multimodal field data, I utilized an environmental and acoustic dataset collected from inside a honeybee (*Apis mellifera*) hive. A custom-built, low-power data logger controlled by an Arduino Pro Mini microcontroller (3.3V, 8MHz; SparkFun Electronics, USA) was deployed within a hive located in Tsukuba, Japan (36.0209° N, 140.1202° E; Fig. 7).

**Fig. 7:**
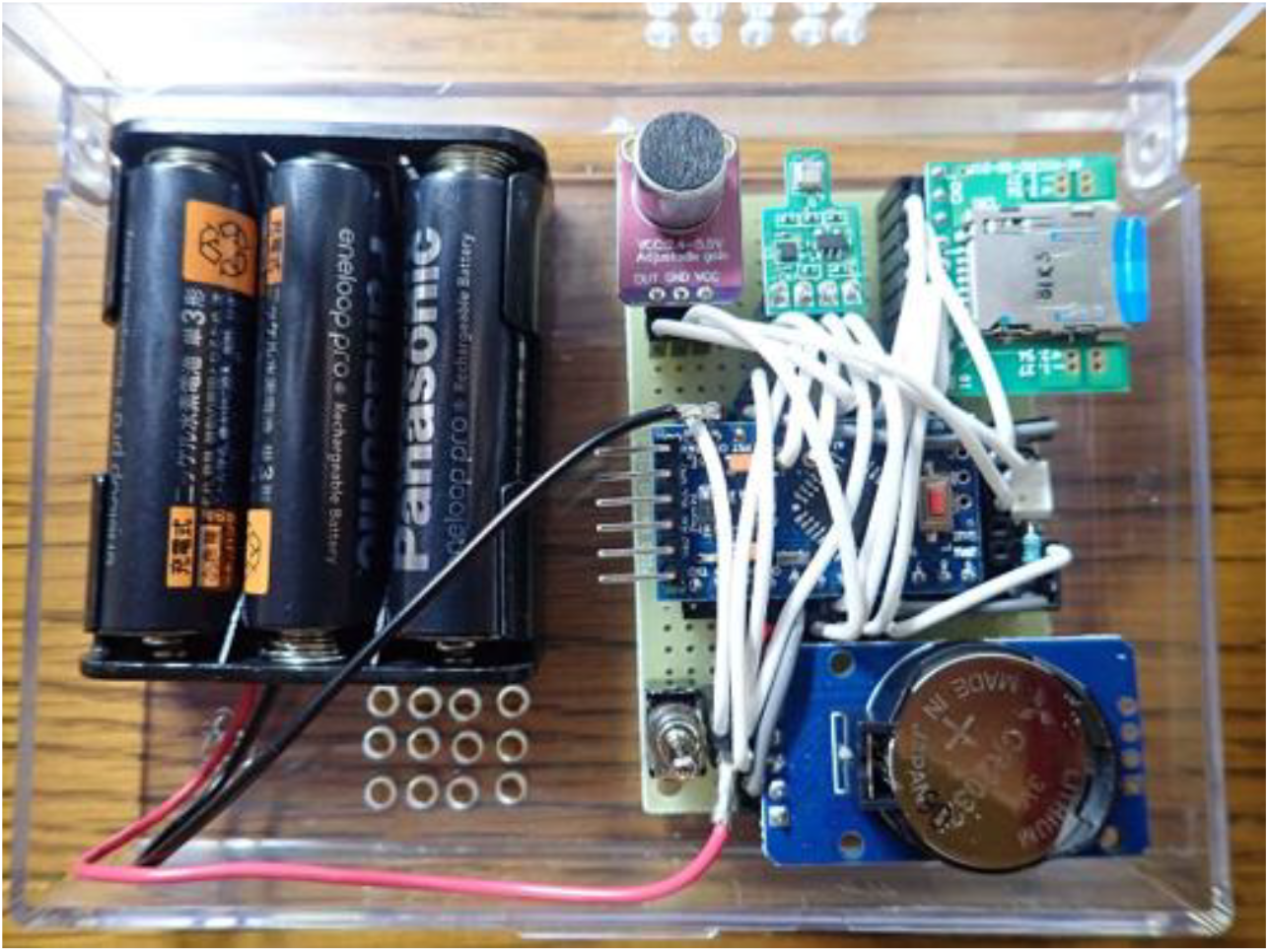
A custom-built data logger placed for data monitoring in a honeybee hive.

Internal temperature was measured using a BME680 environmental sensor module (Bosch Sensortec, Germany), while internal hive sounds were simultaneously captured using a MAX4466 microphone module (Maxim Integrated, USA). Data for both modalities were recorded at 15-minute intervals. During each interval, the audio signal was sampled at a rate of approximately 4,274 Hz to collect 50,000 data points, resulting in an 11.7-second continuous recording. This monitoring was conducted over a period from May 20 to 22, 2022.

Raw audio data were first converted into standard WAV format, and divided into 2-second segments. For each segment, a spectrogram was generated using a Short-Time Fourier Transform (STFT) with a window size of 512 and a 75% overlap. Then each 2-second spectrogram was compressed into a 10-by-10 grid, comprising 10 bins along the time axis and 10 bins along the frequency axis, by calculating the median value within each bin. The pooled 10 × 10 matrices from the segments were then aggregated to create a representative spectrogram for each interval, which was subsequently flattened into a 100-dimensional acoustic feature vector. Finally, each dimensional vector was normalized.

Using the processed multimodal dataset, I applied FORWC to infer the directed causal interactions between the internal hive temperature (a 1D scalar variable) and the acoustic dynamics (a 100-dimensional vector). To evaluate the causality from sound to temperature, I first performed a dimensional screening across a time lag range of *tp* ∈ [–3, 3]. This procedure identified a subset of 49 acoustic time series that possessed significant causal information regarding temperature. For this 49-dimensional acoustic subset, the optimal embedding dimension was determined to be *E*\* = 2, and the optimal time lag was *tp* = –3. Because the target (causal) variable was multidimensional, I executed FORWC using a multi-regression forest to simultaneously cross-map these 49 acoustic states from the temperature manifold. Conversely, to assess the reverse causality—from temperature to sound—I performed a similar screening and identified 61 acoustic dimensions that were significantly affected by temperature. In this direction, where the temperature time series served as the causal target, the optimal embedding dimension was *E*\* = 2, and the optimal time lag was *tp* = 0.

### Computing Environments

All calculations and analyses were performed on the free statistical software R version 4.4 (R Core Team, 2026). To carry out S-map and CCM, ‘rEDM’ (https://github.com/ha0ye/rEDM) was used. For MDR S-map, ‘macam’ version 0.2.5 (https://github.com/ong8181/macam) and rUIC version 0.9.15 (https://github.com/yutakaos/rUIC) were used. To calculate forest weights, ‘grf’ package (Tibshirani et al., 2026) was used.

## Artificial Intelligence

Google Gemini was utilized as a coding assistant to refine, parallelize, and optimize the R scripts for data analysis and visualization. The author completely verified and evaluated all codes to ensure computational accuracy and reproducibility.

## Acknowledgments

I thank Dr. Masayuki Ushio, Dr. Chih-hao Hsieh, and Dr. Takehiko Yamanaka for invaluable discussions and comments on the previous draft. During the preparation of this manuscript, the author used an AI assistant (Google Gemini) for editing and proofreading to improve the readability of the English text. After using this tool, the author thoroughly reviewed and edited the content and takes full responsibility for the final publication.

## Author contributions

S.K. solely worked on conceptualization, methodology, investigation, visualization, supervision, writing, reviewing of this study.

## Data, code, and materials availability

The simulation datasets, experimental time series, and empirical honeybee monitoring data analyzed in this study are available from the corresponding author upon reasonable request. The underlying R scripts and implementation algorithms are subject to institutional intellectual property management related to a pending patent application (Japanese Patent Application No. 2026–87255) and will be made available upon reasonable request for non-commercial academic research and reproducibility verification purposes.

## REFERENCES

Athey, S., Tibshirani, J., & Wager, S. (2019). Generalized random forests. Annals of Statistics, 47(2), 1148–1178.

Breiman, L. (2001). Random forests. Machine Learning, 45(1), 5–32.

Chang, C.-W., Miki, T., Ushio, M., Ke, P.-J., Lu, H.-P., Shiah, F.-K., & Hsieh, C.-H. (2021). Reconstructing large interaction networks from empirical time series data. Ecology Letters, 24(12), 2763–2774.

Ebisuzaki, W. (1997). A method to estimate the statistical significance of a correlation when the data are serially correlated. Journal of Climate, 10(9), 2147–2153.

Hastie, T., Tibshirani, R., & Friedman, J. (2009). The elements of statistical learning: Data mining, inference, and prediction, second edition (2nd ed.) [PDF]. Springer.

Jost, C., & Ellner, S. P. (2000). Testing for predator dependence in predator-prey dynamics: a non-parametric approach. Proceedings of the Royal Society of London. Series B: Biological Sciences, 267(1453), 1611–1620.

Kawatsu, K. (2024). Local-manifold-distance-based regression: an estimation method for quantifying dynamic biological interactions with empirical time series. Royal Society Open Science, 11(7), 231795.

Lin, Y., & Jeon, Y. (2006). Random forests and adaptive nearest neighbors. Journal of the American Statistical Association, 101(474), 578–590.

Munch, S. B., Rogers, T. L., & Sugihara, G. (2023). Recent developments in empirical dynamic modelling. Methods in Ecology and Evolution, 14(3), 732–745.

R Core Team. (2026). R: A language and environment for statistical computing. R Foundation for Statistical Computing.

Sugihara, G. (1994). Nonlinear forecasting for the classification of natural time series. Philosophical Transactions. Physical Sciences and Engineering, 348(1688), 477–495.

Sugihara, G., May, R., Ye, H., Hsieh, C.-H., Deyle, E., Fogarty, M., & Munch, S. (2012). Detecting causality in complex ecosystems. Science (New York, N.Y.), 338(6106), 496–500.

Takens, F. (1981). Detecting strange attractors in turbulence. In Lecture Notes in Mathematics (pp. 366–381). Springer Berlin Heidelberg.

Tibshirani, J., Athey, S., Friedberg, R., Hadad, V., Hirshberg, D., Miner, L., Sverdrup, E., Wager, S., & Wright, M. (2026). Generalized random forests. CRAN.

Veilleux, B. G. (1979). An analysis of the predatory interaction between paramecium and didinium. The Journal of Animal Ecology, 48(3), 787.

Wager, S., & Athey, S. (2018). Estimation and inference of heterogeneous treatment effects using random forests. Journal of the American Statistical Association, 113(523), 1228–1242.

Ye, H., & Sugihara, G. (2016). Information leverage in interconnected ecosystems: Overcoming the curse of dimensionality. Science (New York, N.Y.), 353(6302), 922–925.

